# A novel computational framework to identify translational potential of genetic mouse models in rare genetic obesity

**DOI:** 10.1101/2025.04.22.649931

**Authors:** Nathalie Barreto Lefevre, Aleksandra Polosukhina, Ana Catarina Barata, Michel Leibovici, Pamela Le Quéré, Marcelo Paez-Pereda, Soham Saha, Franck Oury

**Author notes:** These authors contributed equally to this work (Nathalie Barreto Lefevre, Aleksandra Polosukhina).

## Abstract

Finding the right animal models has been a critical bottleneck in translational sciences of rare diseases. Traditional methods can be biased, limiting their effectiveness, and do not include cross-analysis with human manifestations of the diseases. This research introduces a novel computational framework that analyzes hundreds of mouse model data and phenotypes simultaneously. This paves the way to make the best disease-specific decisions. We have identified 106 mouse models in genetic obesity with more than 600 different phenotypes. An evaluation of the shared phenotypes across these models unearthed unexpected connections between obesity and 16 other diseases, opening doors for entirely new treatment avenues. This analytical framework has the potential to modernize how we research rare diseases, leading to more effective therapies for a wider range of conditions.

**Author Summary:** A novel computational approach pinpoints ideal mouse models for rare diseases, uncovering hidden connections with other conditions.

## Introduction

The use of animal models in preclinical and translational research relies on the replication of pathophysiological conditions found in human patients, at different stages of disease progression. An ideal animal model of a human disease is characterized by the similarities they share with human conditions, in terms of pathophysiology of disease (1), phenotypical and histopathological characteristics (2), predictive biomarkers for course or prognosis (3), response to therapies (3, 4), and drug safety and/or toxicity (2). Animal model studies not only provide key insights into mechanism of the disease, but also pave the way to the first trials in humans in terms of dosage determination, expected clinical outcomes, efficacy, toxicity, and safety (5, 6).

Animal models have greatly facilitated the understanding of disease pathology and progression (7–9). This is primarily due to the evolving methods that enable the selection of animals (mice, rats, hamsters, zebrafish, etc.) with specific and variable genetic backgrounds, which mimic vital signs of disease and provide the opportunity to manipulate their genome to obtain desired outcomes in disease modeling (9). These translational models also explore how genes and environmental conditions interact to drive disease progression (10). Additionally, they help identify genetic factors affecting treatment response and pave the way for breakthrough, first-in-class, and best-in-class therapies.

Rare diseases are characterized by low prevalence (1/2000 people in European Union) and present a wide variety of symptoms. A major difficulty in understanding rare diseases stems from the lack of relevant mouse models. A crucial challenge is the identification of existing genetic animal models for a particular disease in an unbiased manner and determining which ones best capture the phenotypes of the disease (11). As data in rare diseases is scarce and fragmented, it’s difficult to determine the direct translational impact in terms of disease progression and therapy development. The identification of overlapping phenotypes across animal models presents an opportunity to map disease adjacencies. This knowledge can be leveraged by researchers to optimize resource allocation during drug discovery and development (12).

Our goal is to find patterns across animal models of different diseases, revealing new links between the human diseases they represent. This approach, using phenotypic similarity to group models of distinct diseases, challenges traditional disease classifications. Our study addresses these key points and harmonizes differences among animal model data using a data-science and human-in-the-loop approach. We designed an automated scoring matrix to account for relevant and repetitive phenotypes across different mouse models. This work facilitates the identification of highly penetrant mouse models, characterized by phenotypes frequently observed in preclinical studies. To bridge the gap between mouse model phenotypes and their human disease counterparts, we have developed an ancillary computational framework leveraging Natural Language Processing (NLP). This framework enables the alignment of mouse model phenotypes with human clinical features documented within the Human Phenotype Ontology (HPO) (13). Additionally, we employed a similar NLP approach to extract and match these phenotypes with established clinical trial endpoints.

We focused on genetic obesity in mice, a rare disease which presents different physiological manifestations. We analyzed 106 models with diverse phenotypes, automatically assessing the frequency of each phenotype and its potential for drug development (translational potential). We identified several clinically relevant phenotypes that could be incorporated as future human clinical trial endpoints (14, 15). Incidentally, we observed that some of the phenotypes represent neuronal features, providing interesting insights into multi- systemic disorders such as obesity. Furthermore, by analyzing shared phenotypic features across models, we established “disease adjacencies” within obesity, allowing for more efficient management of the disease throughout its lifecycle.

## Results

### Selection of optimal mouse models for disease of interest

We have designed an unbiased and curated computational framework to address key constraints in translational rare disease research (Fig 1). Leveraging the power of data science, natural language processing (NLP) and machine learning, we have aggregated information on over 6500 genetic mouse models reported for 1796 rare diseases. In addition, we have attributed the phenotypes exhibited by these mouse models (n = 7685) to 28 organ systems. The present database records over 1900 genes that are reported to be perturbed and/or mutated in these mouse models, with 6427 allelic compositions. The objective was to determine granular, scientific and translational intelligence in a disease of choice. While investigating these mouse models for the level of granularity, we observed that only 8% of the obese mice displayed neurological features.

**Fig 1.**
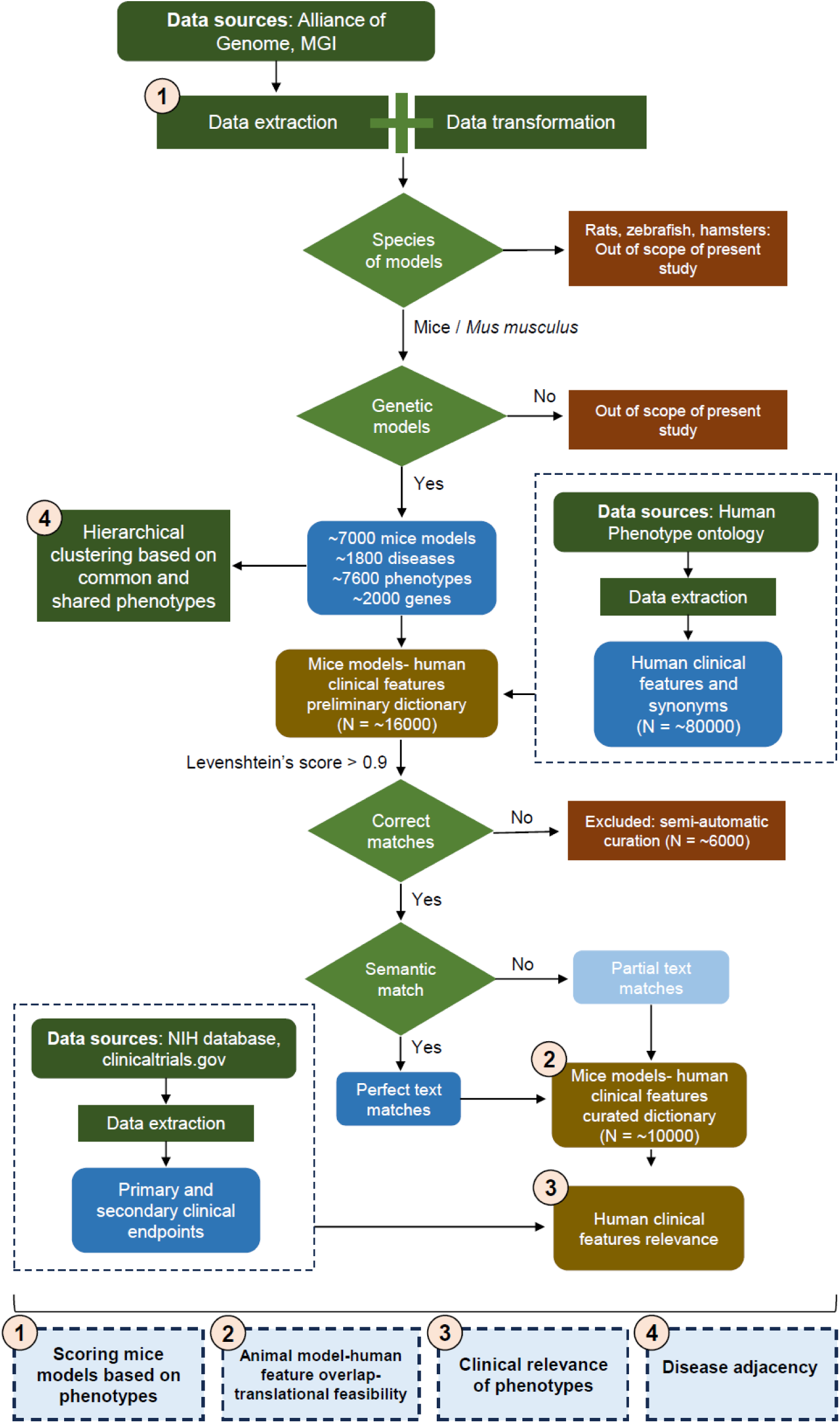
Schematic representation of the workflow of the genetic mice model framework. The key functionalities include scoring the mouse models for relevance based on phenotypic descriptions, translational feasibility based on mouse model-human feature overlap, clinical relevance, and disease adjacency.

We identified 106 mouse models described as having genetic obesity, with 596 reported phenotypes across 28 organ systems. 17 genes were reported in those mouse models with 93 allelic compositions. The scoring of different mouse models with the reported phenotypes demonstrates a wide variety of phenotypic descriptions (Fig 2A, S1 Table). The key phenotypes with the highest frequency across the models were obese (n=66), increased circulating insulin (n=47), increased leptin (n=31), hyperglycemia (n= 29), increased body weight (n=27), insulin resistance (n=24), increased triglycerides (n=23), increased cholesterol (n=23), polyphagia (n=22), impaired glucose tolerance (n=21), increased liver triglyceride (n=20) and increased liver weight (n=16) (Fig 2B). Among the systems most impacted by the phenotypes in the mouse models were metabolism (37%), growth (15%), and adipose tissue (9%) (Fig 2C, S1 Table). Interestingly, neurological/behavioral phenotypes account for 8% of all the observed phenotypes while reproductive system was implicated in 6% (Fig 2C, S1 Table). Among the diseases represented by the models, 61% were assigned to general obesity, 22% to Prader-Willi syndrome, 16% to Abdominal obesity-metabolic syndromes, and 1% to Cohen syndrome (Fig 2D).

**Fig 2.**
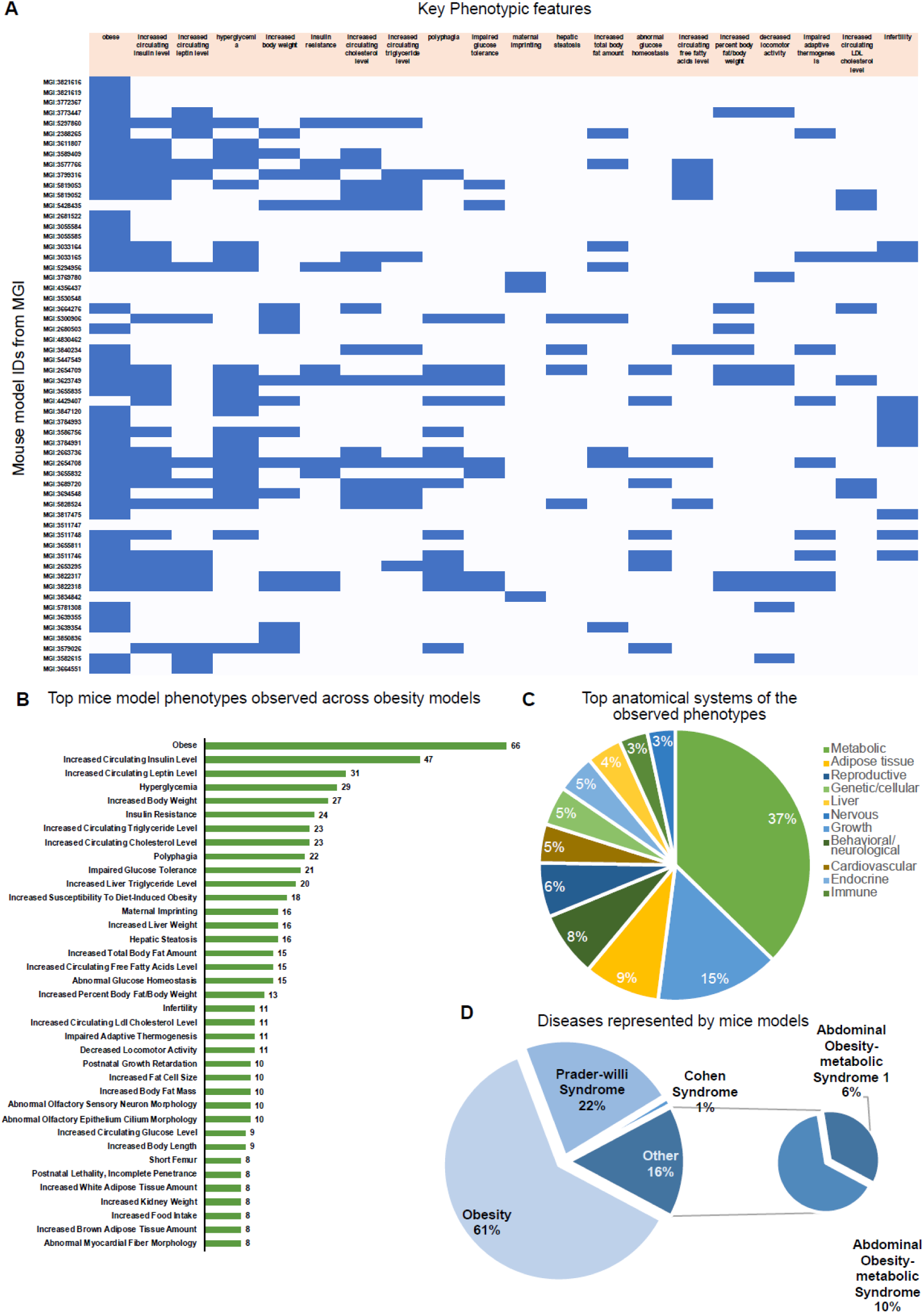
Demonstration of the scoring matrix and intelligence generation from mouse models of obesity. **(A)** An extract of the scoring matrix with respect to phenotypes that are relevant for obesity. Note that some phenotypes are repeated across different mouse models. Blue indicates the presence of the phenotypes while white indicates that the phenotype is absent or not reported in the model. **(B)** Bar plot showing the counts of the top phenotypes observed across different mouse models of obesity. **(C)** Distribution of the key systems encompassed by the mouse model phenotypic descriptions. Note that metabolic system represents the highest portion of the phenotypes. **(D)** Distribution of the mouse models describing diseases with obesity as a disease or a clinical feature.

### A computational framework for selecting mouse models replicating human features

A key determinant in translational feasibility of drug design and development is the strength and degree of mouse models to represent human disease conditions. In order to facilitate the final translation of biomarkers from basic research into human applications, animal models need to be improved to yield more translatable results (*16*). Our framework, designed to link human clinical features to observed mouse phenotypes, addresses one of the key points in translational studies: similarity determination between mouse phenotypes and human symptoms.

In our study on obesity, we matched mouse phenotypic ontologies reported in Mouse Genome Informatics (MGI) to human phenotype ontologies reported in Human Phenotype Ontology (HPO). We devised two distinct frameworks of phenotype matching: a perfect match, where text from either component had a 100% lexical equivalence (example, “abnormal eye physiology” in Fig 3A, S2 Table), and a partial match, where the text did not have 100% lexical equivalence but demonstrated semantic similarity (example, “impaired hearing” matched to “mixed hearing impairment” in Fig 3A, S2 Table). The overall percentage of matches was calculated taking both these lexical matches into account. The percentage total of lexical matches between mouse phenotypes and human clinical features ranged from 66.67% in certain models to 20% (Fig 3B, range shown for models with 51.22%-66.67% matches). In our semi-automatic workflow, around 6000 matches were manually curated and excluded based on the degree of semantic sense (Fig 1). Detailed meta-analysis in the term matches indicated that the proportion with the highest term matches were “increased body weight” (10%), “tall stature” (10%), “hyperglycemia” (8%), “hypertension” (8%), “insulin resistance” (7%), “glucose intolerance” (6%) and others (Fig 3C).

**Fig 3.**
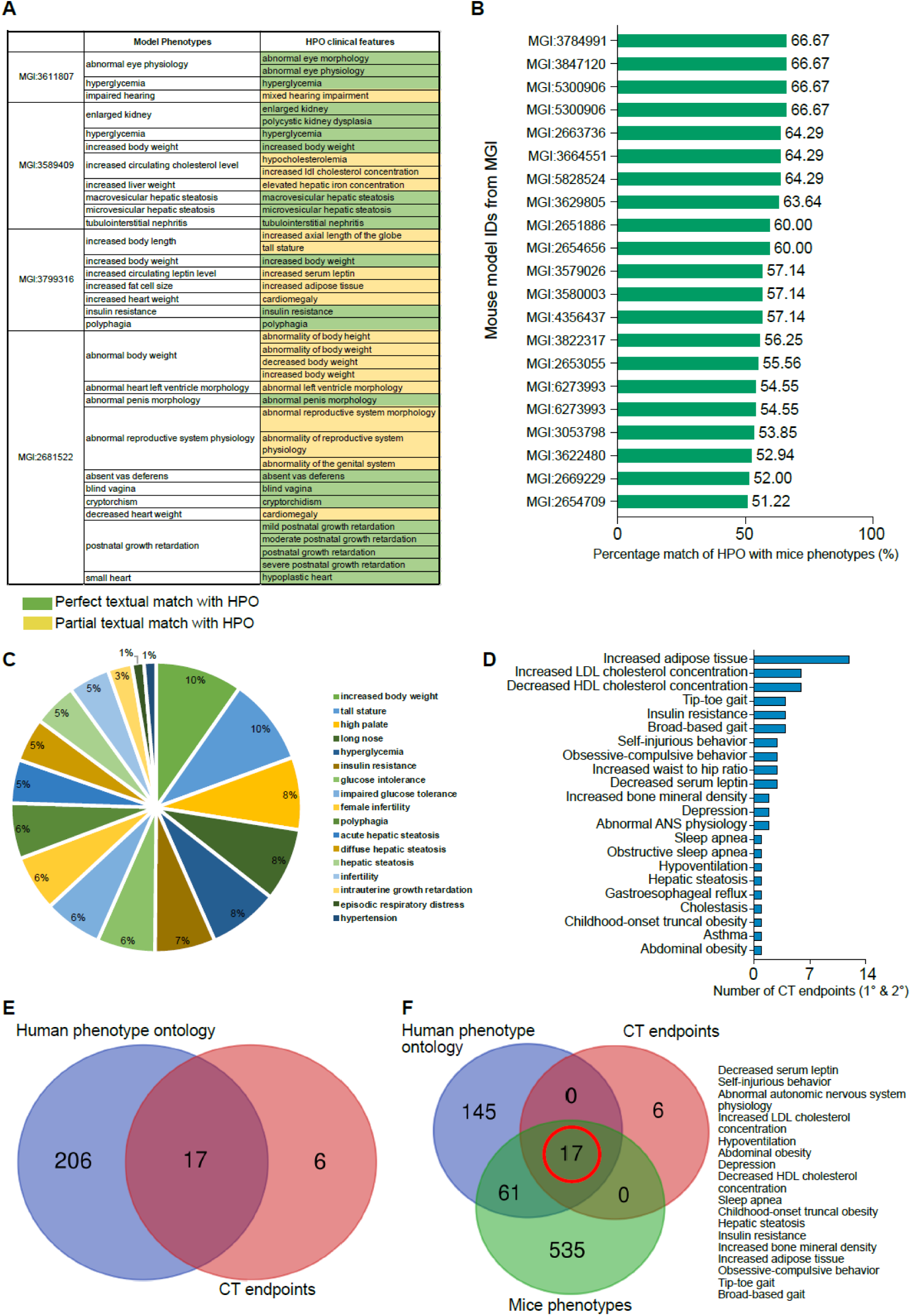
Phenotype matching of mouse models with human clinical features. **(A)** An extract of the matching between mouse model phenotypes and human phenotypic descriptors from HPO. The green color indicates a perfect semantic match while the yellow indicates a partial semantic match. **(B)** Quantification of the percentage matches of mouse model phenotypes that match with HPO phenotypes. The percentage indicates total number of matches observed (partial and perfect). **(C)** Distribution of the key semantic matches across mouse models. **(D)** Bar plots indicating the frequency of the primary and secondary (1° & 2°) clinical endpoints matched to the HPO phenotypes. **(E)** Overlap analysis of all HPO phenotypes with relevant clinical endpoints indicating 17/23 clinical endpoints could be semantically matched to relevant HPO phenotypes. **(F)** Overlap analysis of HPO phenotypes, mouse model phenotypes and clinical trial endpoints indicate that 17 phenotypes matched in panel E had a corresponding clinical trial endpoint in all obesity trials.

### A computational framework for selecting clinically relevant model systems

Determining the clinical relevance of mouse models requires validation of the model’s accuracy in reproducing human disease pathogenesis and response to treatment. Our objective was to design an efficient way to assign clinical relevance to the observed mouse phenotypes that matched with human disease phenotypes.

In clinical trials, researchers assess the effects of a new treatment by measuring specific outcomes. Primary endpoints are the main outcomes that the trial is designed to measure. They are the most important factors in determining whether the new treatment is effective. Secondary endpoints are additional outcomes that can provide supporting evidence about the effectiveness of the new treatment, or they can explore other potential benefits or risks associated with the treatment. Choosing the right endpoints is crucial for the success of a clinical trial and estimating the amount of overlap of these endpoints with mouse models is critical to evaluate the translational potential of any drug. We extracted the primary and secondary clinical trial endpoints of the reported clinical trials in each disease and adapted the previous framework to match terms in clinical trials to matched-HPO phenotypes. A key challenge in this framework is the lack of a consistent dictionary in endpoints which make the matching comparatively more difficult. Our framework revealed that the most frequent clinical endpoint across obesity was increased adipose tissue (n=12), decreased High-Density Lipoprotein (HDL) cholesterol concentration (n=7), increased Low-Density Lipoprotein (LDL) cholesterol concentration (n=7), insulin resistance (n=4), and increased waist to hip ratio (n=3) (Fig 3D, S3 Table). Neurological and behavioral observations were also implicated such as self-injurious behavior (n=3), obsessive compulsive behavior (n=3), abnormal autonomic nervous system (n=2), and depression (n=2) (Fig 3D, S3 Table).

Comparing all the reported clinical endpoints to the HPO phenotypes, we observed that 17 out of 23 uniquely reported clinical endpoints matched relevant clinical phenotypes in humans (Fig 3E, S3 Table). Some of these matched features include decreased serum albumin, increased LDL cholesterol, abdominal obesity, decreased HDL concentration, hypoventilation, sleep apnea, depression, and others. Expanding the overlap with mouse model phenotypes, we found that the 17 matched clinical endpoints were already replicated in mice (Fig 3F). Interestingly, we observed almost 400 new features in mice that are matched to human phenotypes (such as cardiomyocyte hypertrophy, abnormal liver physiology, hepatic steatosis and enlarged kidney), enhancing our repertoire of addressable clinical features that could be replicated in a non-human preclinical model.

### Identification of disease adjacencies

In comparative disease modeling, exploiting phenotypic similarities across animal models offers a powerful approach. Shared phenotypes can unveil conserved pathophysiological mechanisms, potentially translating preclinical findings to humans.

The present case study with obesity resulted in identifying 16 unique disease models (Fig 4A, S4 Table), segregated across 4 distinct clusters. The diseases include adrenal gland disease, Bardet-Biedl syndrome, disease of metabolism, endocrine system disease, primary hyperaldosteronism, acquired metabolic disease, Prader-Willi syndrome, Alstrom syndrome, obesity, Type 2 diabetes mellitus, disorder of sexual development, maturity-onset diabetes of the young, autoimmune disease of endocrine system, primary immunodeficiency disease, Albright’s hereditary osteodystrophy, metal metabolism disorder, and Smith-Magenis syndrome. Among the 4 clusters with obesity as a phenotype, we observed differential features based on the phenotypes in mouse models. The key features that are similar across the clusters include involvement of adipose tissues, growth of body size, reproductive system, behavior and neurology, homeostasis and metabolism. Clusters C1 and C2 are specified by cellular components, while C3 diseases are involved in the cardiovascular system, endocrine exocrine glands, and the liver biliary system (Fig 4B, S4 Table). Cluster C4 demonstrates neurological phenotypes (Fig 4B). To validate the adjacency of the diseases obtained in the previous observation, we mapped the reported genes against the diseases and analyzed the gene networks (Fig 4C). Gene network and enrichment analysis identified key processes linked to obesity, including adipose tissue development, lipid metabolism, adipogenesis, differentiation of white and brown adipocyte, Prader-Willi and Angelman syndromes, and neurological manifestations (Fig 4D, S5 Table). In terms of phenotypes over-represented by the genes, processes linked to abnormal circulating inhibin levels, bulimia, central adrenal insufficiency, hypothyroidism, hypopituitarism, abnormal T4 concentration and temperature sensation were enriched (Fig 4D-E). The identified over-represented phenotypes suggest a complex genetic underpinning of obesity beyond metabolic pathways. These genes appear linked to a broader spectrum of physiological processes, including hormonal regulation, appetite control, and sensory perception. The association with conditions like bulimia, thyroid dysfunction, and adrenal insufficiency indicates a potential interplay between genetic factors and the development of obesity (Fig 4E, S5 Table). This multifaceted genetic influence highlights the need for a comprehensive approach to understanding and treating obesity.

**Fig 4.**
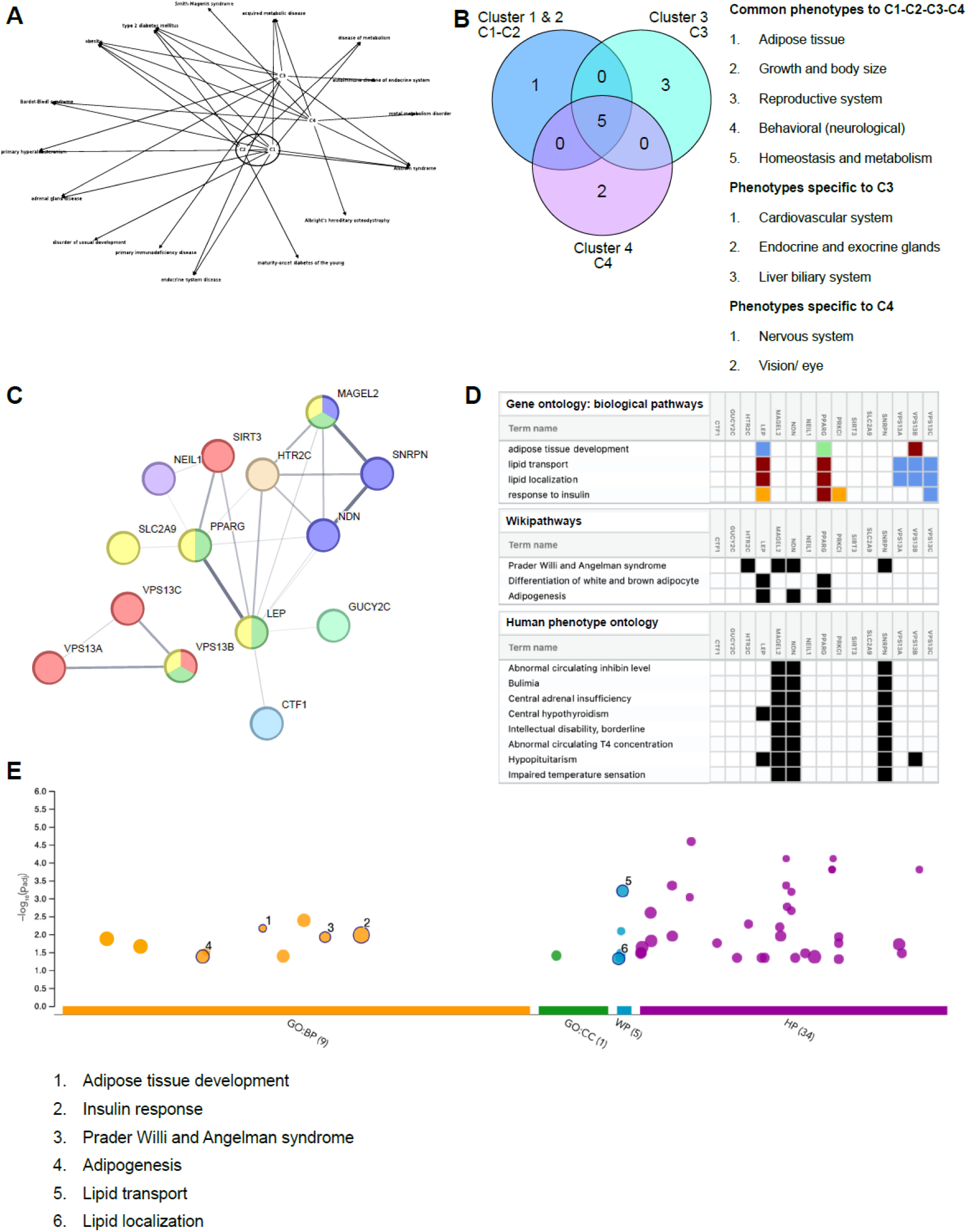
Disease adjacency evaluation. **(A)** A graphical representation of the key diseases with obesity and the corresponding clusters. Cluster C1 and C2 seem closer to each other compared to Cluster C3 and Cluster C4. All clusters share similarities in adipose tissue, growth of body size, neurological involvement and anomalies of metabolism. **(B)** Overlap analysis of the phenotypes in each cluster indicates that C1 and C2 comprise components of cellular nature in obesity, while C3 contains cardiovascular, liver, and endocrine/exocrine components. C4 represents features with neuronal involvement. **(C)** Gene network of the key genes described in Clusters C1, C2, C3, and C4. **(D)** Enrichment analysis in g:profiler indicates that key processes (GO:biological processes, WikiPathways and Human phenotype ontology) represented by these genes are linked to obesity, adipose tissue development, lipid transport and localization, adipogenesis, Prader-Willi and Angelman syndrome, and phenotypes linked to neurological manifestations of obesity, bulimia and thyroid hormone disbalance. **(E)** Graphical representation of enrichment analysis performed in panel D. Top processes include organic substance transport, Prader-Willi and Angelman syndrome, Insulin response, Adipogenesis and Lipid transport and localization. BP, Biological Process; CC, Cellular Component; HP, Human Phenotype; WP, WikiPathways.

## Obesity mouse models with neuronal phenotypes

Finally, to estimate the level of granularity provided by the analytics, we deep dived into the neurological phenotypes of these mouse models. We observed 21 mouse models with different model phenotypes related to the nervous system processes (Fig 5A), five of which (MGI: 3694548, MGI: 2174793, MGI: 4429407, MGI: 3773672 and MGI: 2653055) consisted of the highest number of phenotypes related to the nervous system (Fig 5A, B).

**Fig. 5.**
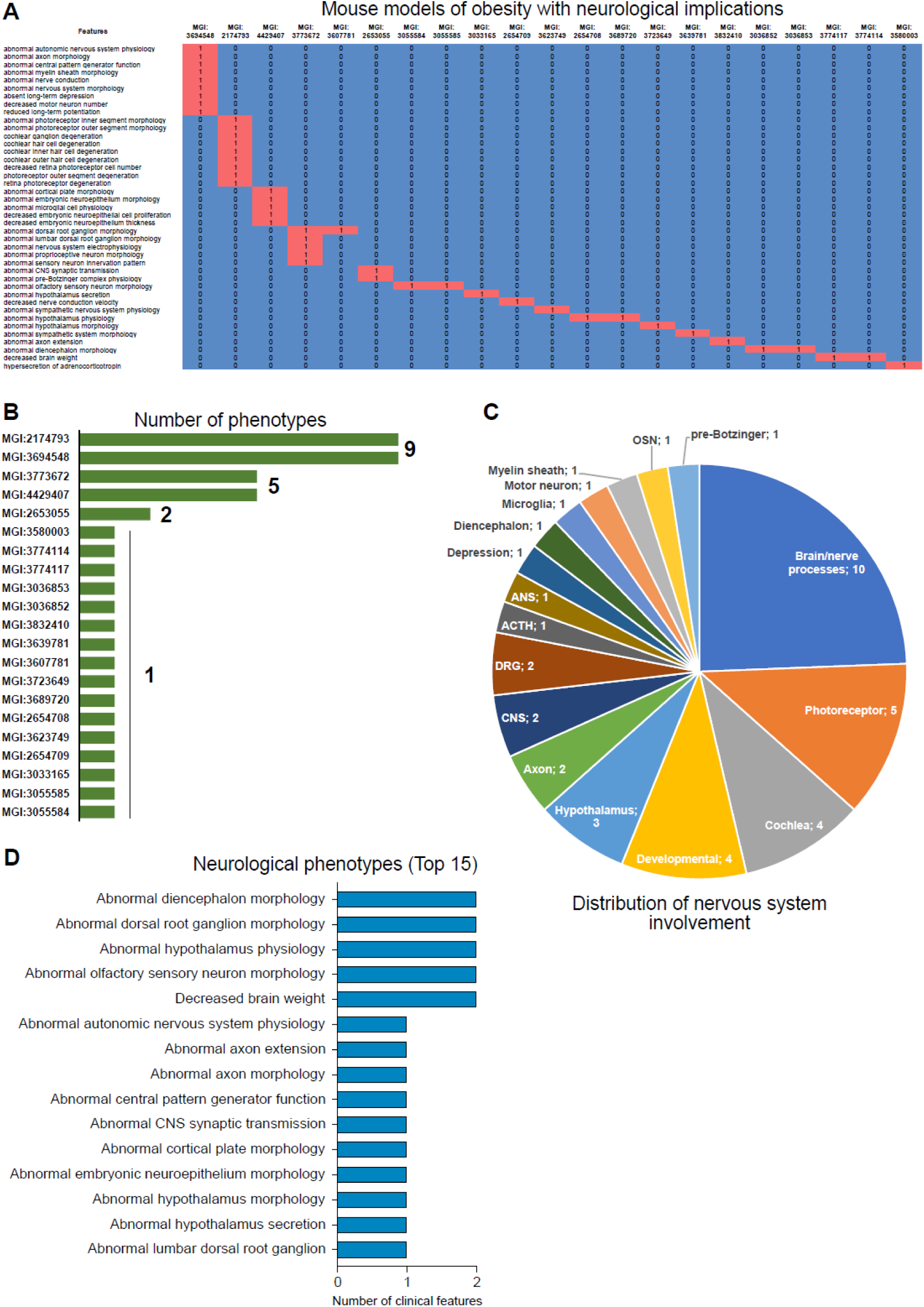
Evaluating obesity mouse models showing neuronal phenotypes. **(A)** An extract of the scoring matrix demonstrating obesity mouse models with neurological impact. **(B)** Quantification of the total number of neurological phenotypes indicate that 5 mouse models have more than one phenotype involving the nervous systems. **(C)** Distribution of key neuronal substrates described by the mouse models of obesity. **(D)** Bar plot showing the top 15 neuronal phenotypes described in the mouse models of obesity.

The detailed model phenotypes described in these models and their corresponding system are summarized below in Table 1:

**Table 1:** Obesity mouse models with neurological implications.

| Model name | Phenotypes observed | Systems / Organs classified |
| --- | --- | --- |
| MGI: 3694548 | Abnormal ANS physiology<br>Abnormal axon morphology<br>Abnormal central pattern<br>Abnormal myelin sheath morphology<br>Abnormal nerve conduction<br>Abnormal CNS morphology<br>Long term depression<br>Decreased motor neurons<br>Reduced LTP | ANS, CNS,<br>Myelin sheath,<br>Depression, Motor<br>neuron, Brain/<br>nerve processes |
| MGI: 2174793 | Abnormal photoreceptor inner segment morphology<br>Abnormal photoreceptor outer segment morphology<br>Cochlear ganglion degeneration<br>Cochlear hair cell degeneration<br>Cochlear inner hair cell degeneration<br>Cochlear outer hair cell degeneration<br>Decreased retina photoreceptor<br>Photoreceptor outer segment degeneration | Photoreceptors,<br>Cochlea |
|  | Retina photoreceptor degeneration |  |
| MGI: 4429407 | Abnormal cortical plate morphology<br>Abnormal embryonic neuroepithelium morphology<br>Abnormal microglia cell physiology<br>Abnormal embryonic neuroepithelium cell proliferation<br>Abnormal embryonic neuroepithelium thickness | Microglia, developmental |
| MGI: 3773672 | Abnormal DRG morphology<br>Abnormal lumbar DRG morphology<br>Abnormal nervous system electrophysiology<br>Abnormal proprioceptive neuron physiology<br>Abnormal sensory neuron innervation pattern | DRG, Brain/ nerve processes |
| MGI: 2653055 | Abnormal CNS synaptic transmission<br>Abnormal pre-Botzinger complex physiology | CNS, pre-Botzinger complex |

Among the key neuronal phenotypes, major processes were observed to be involved in brain/nerve processes (n=10), photoreceptors (n = 5), cochlea (n = 4), developmental processes (n = 4), and the hypothalamus (n = 3) (Fig 5C). Depression like phenotype was seen in one of the mouse models (Fig 5C). Involvement of microglia, motor neuron, diencephalon, myelin sheath, and olfactory sensory neurons (OSNs) appeared as other brain axes implicated among the mouse models for obesity (Fig 5C). To identify commonalities amongst the observed phenotypes, we performed a segregation analysis based on major organs or neuronal systems. This analysis revealed sensory physiology, hypothalamic physiology, dorsal root ganglion (DRG) and diencephalon morphology as the most frequently affected axes (Fig 5D).

## Discussion

There are approximately ∼7000 rare diseases described in the world, including clinical subtypes, anomalies, and morphological alterations (*17*). Most of these diseases have poor scientific understanding with respect to biological mechanisms and disease progression states. This not only poses significant challenges in treatments, but also delays therapy development. In addition, the time constraint to diagnose rare disease patients correctly is also time consuming. Our present work provides a framework to understand rare diseases from the genetic mouse models perspective. This present framework offers new insights into the preclinical similarities across different model systems and tries to fill in the knowledge gaps that exist with the human presentation of these rare diseases. Using the overarching power of advances in data science and analytics, we aimed at harnessing the power of phenotypic data across mouse models and humans, and assess the impact of these phenotypes on the disease (*18*, *19*).

Identifying phenotypic similarities across multiple diseases leads to fast and precise diagnoses of rare diseases and is an important step to stratify patients into meaningful treatment subgroups (*20*). This is particularly difficult in genetic diseases (which accounts for almost 80% of rare diseases) as drawing a significant genotype-phenotype correlation is difficult. This is because a single gene can confer significant phenotypic heterogeneity (*21*). Such initiatives on semantic similarity offer enormous insights into disease phenotypes and help capture the diagnostic information embedded in the phenotype ontology (*22*). Clustering genetic mouse models of rare diseases based on their expressed phenotypes confers the ability to study rare diseases as a group of phenotypically similar phenomenon rather than in isolation. This allows us to design basket and/or bucket trials of groups of rare diseases that can be grouped by virtue of the phenotypes they share. These higher order relationships among diseases could be represented in the context of a network to homogenize data across multiple related diseases.

In this study, we explored the scalability of the proposed framework with a rare group of diseases such as genetic obesity. We have developed high-performance tools allowing for a rapid detection of valuable mouse models replicating human rare diseases. Obesity remains a global health crisis with several unmet medical needs. While there are approved therapies for obesity, they have significant side effects and do not address the whole spectrum of obesity patients, mainly genetically obese patients.

A comprehensive and holistic multidisciplinary approach that leads to a deep biological understanding and addresses the psychological, behavioral, and social determinants of obesity is still lacking in many healthcare systems. Lastly, despite its widespread prevalence, there is a significant gap in tailored preventive strategies and early interventions for at-risk populations, emphasizing the need for personalized and proactive healthcare solutions.

The presented framework tackles these challenges and allows the expansion of rare disease knowledge to greatly increase the scope of translational relevance of preclinical research. It is designed to address rare disease knowledge gaps to efficiently identify new uses for existing drugs (drug repurposing), discover relevant biomarkers, and design efficient basket trials across phenotypically similar diseases.

In our example of genetic mouse models linked to obesity, we have validated some of the key observations that have been previously described. Among them are features of adipose tissue deposition (*23*), hormonal imbalances (*24*), and cholesterol alterations (*25*). Some of these key recapitulative features have been observed in humans (*26*) as clinical trial endpoints.

By leveraging phenotypic data and their corresponding gene networks, we can improve diagnoses, patient stratification for targeted therapies, and overall efficiency of drug development for rare diseases. Finally, focusing on shared phenotypes in rare disease models refines research questions, minimizes unnecessary animal testing, and improves overall efficiency.

## Materials and Methods

### Data sources

The study was conducted on open-source data available from different sources. Key data sources used in the study include Alliance of genome (alliancegenome.org) for the list of key genetic animal models, genes and mutations in the animal models and the corresponding phenotypes. The data extracted was cross referenced with the database, MGI-Mouse Genome Informatics-The international database resource for the laboratory mouse (jax.org) to obtain the data related to the nature of the mutation (knock-in, knock-out, transgenic, etc.). For associated human phenotypes in the corresponding diseases, we obtained the clinical features and associated organ systems from the Human Phenotype Ontology (HPO) database (www.hpo.jax.org). Finally, for the clinically relevant features, the primary and secondary endpoints for trials in diseases were extracted from the NIH- database on clinical trials: clinicaltrials.gov.

### Data extraction

We extracted key terms of interest from different vocabularies, such as MGI-Mouse Genome Informatics and Human Phenotype Ontology. Our data extraction concentrated on the following preclinical and clinical entry points- biomarkers, animal model types, key phenotypic features, disease features and key genes. To enrich the extracted information, clinical endpoints (primary and secondary) have been extracted in parallel from clinicaltrials.gov to identify relevant clinical features that can be mapped to mouse phenotypes described in the disease models.

### Data preprocessing

Given the nature of the data used in this project, which includes phenotypes of mouse models, represented through sequences of words, the One-Hot Encoding approach, which is one the most common techniques, is a suitable choice for converting the data into numerical format. Since our phenotypes are diverse and are expressed through a wide vocabulary, converting them into individual binary features helps maintain their uniqueness and allows us to capture the complex relationships between mouse models and their diverse phenotypes without introducing any unintended hierarchical associations between them.

Each mouse model contains a list of phenotypes referring to the observable characteristics, such as physical traits, that the mice exhibit, and which embodies the manifestation of genetic manipulations and environmental interactions.

As the genetic makeup of mice is typically altered to mimic human disease or replicate specific genetic mutations seen in humans, their phenotypic attributes are measured and studied to evaluate the impact of these genetic modifications, to better understand human disease. For each of the 6500+ mouse models, we extracted the associated list of phenotypes, obtaining a total of 7685 unique phenotypes.

For every model, only the binary variables associated to its phenotypes will be "on", representing the affiliation of the model with a particular phenotypic category. For that reason, the initial textual data is converted into numerical data in the format of a matrix with dimension 6500 X 7685, where each row represents a mouse model, and each column corresponds to a phenotypic description. If the model contains the phenotype, its value is 1, otherwise, it will be 0.

The encoding strategy results in a sparse matrix, in which most of the entries are set to 0. The number of phenotypes in each mouse model ranges between [1, 100], with an average of 12 phenotypes per model. This leads to an encoding with an average percentage of zeros in each row equal to 99,5%, reflecting the distinct combination of phenotypic attributes of each mouse model as well as the biological diversity inherent to the phenotypic descriptions of the mouse models.

### Human to mouse traits matching

Human clinical features are described using a standardized vocabulary that depicts phenotypic abnormalities encountered in human diseases - the HPO. Mice also showcase phenotypic traits associated with genetic modifications to mimic human disease; however, the phenotypes aren’t described using the same standardized HPO vocabulary nor do they follow a mouse-specific vocabulary. We created a dictionary to match the human clinical features with the mouse phenotypes to be able to easily identify relevant phenotypes that can be used to track disease progression, at the clinical trial phase. The dictionary associates, for each human clinical feature, the corresponding equivalent mouse phenotype(s), as they often express the same meaning using words that are not necessarily the same. To bridge the gap between the two, we use the Levenshtein distance to evaluate the similarity between the phenotypes and the clinical features. The Levenshtein distance is a text similarity measure (*26*) that compares two words and returns a numeric value representing the total number of single-character edits required to transform one word into another (the distance between two strings). If the Levenshtein distance is equal to zero, the two strings are equal. The larger the Levenshtein distance, the more dissimilar two strings are and the higher the number of single-character operations are necessary to transform one string into the other. To develop the dictionary mapping mouse model phenotypes to human clinical features, we implemented a fuzzy string-matching algorithm, in Python, using the fuzzy-wuzzy library, which is built on the Levenshtein distance, to obtain the similarity score between mouse phenotypes and human clinical features. The closer the score is to 100, the more similar the two strings are. We defined a threshold equal to 80, taking into consideration clinical expertise, for which the scores above this threshold were considered a match.

After manually curating the matches, we obtain a dictionary with total of approximately 10000 matches linking human clinical features to their equivalent mouse phenotypes.

### Clinical relevance of the mouse phenotypes

The lack of a consistent dictionary in clinical endpoint reporting and demonstration makes it difficult to drive meaningful data driven insights from trial reports. A key determinant of a translationally and clinically applicable model system is the ability to represent or replicate viable endpoints in clinical trials (primary and secondary). To achieve a global understanding of the clinical implications of animal models and their phenotypes, we developed a computational framework to create a clinical trial endpoint dictionary matched to the HPO clinical features. To account for lexical noise, we adopted a tokenization approach followed by a fuzzy- matching approach, as described in the human to mouse traits matching section. We used Levenshtein distance to evaluate the similarity between the HPO features phenotypes and the clinical endpoints. Finally, we performed a tri-dimensional analysis of the overlap of features with HPO/clinical endpoints and animal model phenotypes for different rare diseases. This allowed us to build a scoring system to determine which models globally have a higher number of features represented in clinically relevant trials against those diseases. Quality checks for the matches so obtained were performed by randomized manual curation of the matches to improve the percentage of matches so obtained. The frequency of the endpoint terms was also calculated for each disease.

### Disease adjacency

Considering the objectives of this study, we will leverage hierarchical clustering using the complete linkage to identify hidden patterns within our dataset, grouping mouse models based on similar phenotypic attributes and gaining insights on the associations between different mouse models. With the One-Hot encoding representation of our data, we leverage the Jaccard similarity, which measures the proportion of shared elements between mouse models, ranging between 0 and 1 (the closer to 1 the more similarity between the two datasets) to obtain the distance matrix.

Since we do not know the optimal number of clusters beforehand, we use internal cluster validation measures, including the silhouette score, to determine the appropriate number of clusters. The disease heterogeneity within each anatomical system and the uniqueness of the associated mouse models result in a very dense dendrogram, from which we could not extract useful information. Therefore, we set different values for the distance threshold and analyzed the silhouette score while balancing the number of generated clusters.

Low values of the distance threshold result in a number of clusters close to the total number of data points. This indicates that the overall distance between data points is high, reflecting the unique phenotypic profiles of the mouse models and their biological diversity. An elevated number of clusters does not provide insightful information since each cluster would contain very few mouse models.

The objective is also to generate clusters with an acceptable degree of compactness while simultaneously maintaining clear boundaries. In addition to the silhouette score, we leverage the average intra-cluster variation which provides a measure of the average distance between data points within the same cluster, reflecting how close (similar) the data points are to each other. Since we are handling high-dimensional data, the density of data points tends to decrease and the distance between them tends to increase. This can lead the silhouette score to be close to zero as data points may be very close to the boundaries between two neighboring clusters. By measuring the average distance between points within the same cluster, we evaluate the cohesion of the clusters, even if they are not well separated within the feature space. A lower value of the average intra-cluster variation reflects that the points within each cluster are close to each other and an overall compact solution.

After analyzing the evolution of both validation scores, we select the distance threshold that generates a total of 400 clusters.

### Gene network analysis

To confirm the biological relevance of the identified disease clusters, we performed an enrichment analysis and mapping of interactome networks using the genes described for each disease. We used String-db (https://string-db.org) to visualize the gene interactions using full network set up, with network edges defined by confidence of the highest threshold (90%). Gene set enrichment for processes were performed with “g:profiler” (*27*, *28*), with Gene Ontology: Biological processes, Wiki-pathways and Human phenotype ontology dictionaries selected for representation and analysis. Genes represented by the diseases in mice models were mapped to key pathways with a filter of false discovery rate (FDR) set at < 0.05. The p-adj value was taken to be significant at p-adj < 0.05.

## Acknowledgments

We thank Dr. Dimitrije Milunov and Mr. Manish Sarkar from MedInsights for critical reading of the manuscript and feedbacks.

## Author contributions

Conceptualization: AP, NBL, SS, MPP

Methodology: NBL, ACB, SS, AP, PLQ

Investigation: NBL, SS

Visualization: NBL

Supervision: SS, MPP, FO, ACB

Writing—original draft: NBL, SS, ML, FO

Writing—review & editing: SS, ML, FO, ACB, MPP, AP

